# Passive Novel Isodamping Dynamometer for Mobile Measurement of Plantar Flexor Function

**DOI:** 10.1101/2020.06.15.153015

**Authors:** John F. Drazan, Todd J Hullfish, Josh R. Baxter

**Affiliations:** Department of Orthopaedic Surgery, University of Pennsylvania, Philadelphia, Pennsylvania, USA

**Keywords:** plantar flexor, biomechanics, dynamometer, ankle, personalized medicine

## Abstract

Isokinetic dynamometers are the gold standard tools used to assess *in vivo* joint and muscle function in human subjects, however, the large size and high cost of these devices prevents their widespread use outside of biomechanics lab. In this study, we developed a mobile dynamometer to allow for field measurements of joint level function. To ensure subject safety, we designed a new “isodamping” dynamometer that acted as passive energy sink which constrains velocity by forcing incompressible oil through an orifice with an adjustable diameter. We validated the performance of this device by testing plantar flexor function in six healthy adults on both a commercial isokinetic dynamometer and this novel device at three velocities/damper settings and at three different effort levels. During maximal effort contraction, measurements of peak moment and velocity at peak moment of the novel device and the commercial device were strongly correlated along the predicted quadratic line (R^2^ > 0.708, *p* ≤ 0.008). The setting of the damper prescribed the relationship between peak moment and velocity at peak moment across all subjects and effort levels (R^2^ > 0.91, *p* < 0.001). The novel device was significantly smaller (0.75 m^2^ footprint), lighter (30 kg), and lower cost (~$2,200 US) than commercial devices compared to commercially-available isokinetic dynamometers (5.95 m^2^ footprint, 450 kg, and ~$40,000 US respectively).

## INTRODUCTION

Isokinetic dynamometers are the gold standard tools used to assess *in vivo* joint and muscle function in human subjects (Stark et al., 2011). These devices accurately measure the angular position, angular velocity, and the moment generated about a joint. They can also prescribe the range of motion tested and the velocities at which a joint can rotate (Valovich-mcLeod et al., 2004). However, commercial isokinetic dynamometers are expensive and immobile, which limits their use to laboratories and some rehabilitation clinics (Mavroidis et al., 2005). As a result, these types of data cannot be collected in places of interest such as the field or clinic (Verheul et al., 2020), which may explain the high prevalence of small convenience samples (<30 subjects) within human biomechanical research (Vagenas et al., 2018).

Therefore, the purpose of this study was to design and validate a low-cost and mobile dynamometer capable of measuring physiologically relevant joint function outside the laboratory. We chose to use the plantar flexors as a test case due to their importance of ankle kinetics for human locomotion (Neptune et al., 2001) and their utility as a model system to study muscle mechanics (Crouzier et al., 2018). We designed this new device to be portable, low-cost, and safe to use by constraining joint motion with a passive resistance system rather than the active electrical resistance employed in commercial isokinetic dynamometers.

## METHODS

### Device Design

We designed an ‘isodamping’ dynamometer (**Fig. 1A**) to meet several design requirements: 1) use passive components that dissipate energy to ensure subject safety; 2) be low-cost and mobile to facilitate outside-the-lab use. 3) reliably prescribe joint range and rate of motion by adjusting the damper setting; and 4) perform similarly to traditional isokinetic dynamometry when testing human subjects.

**Fig. 1.**
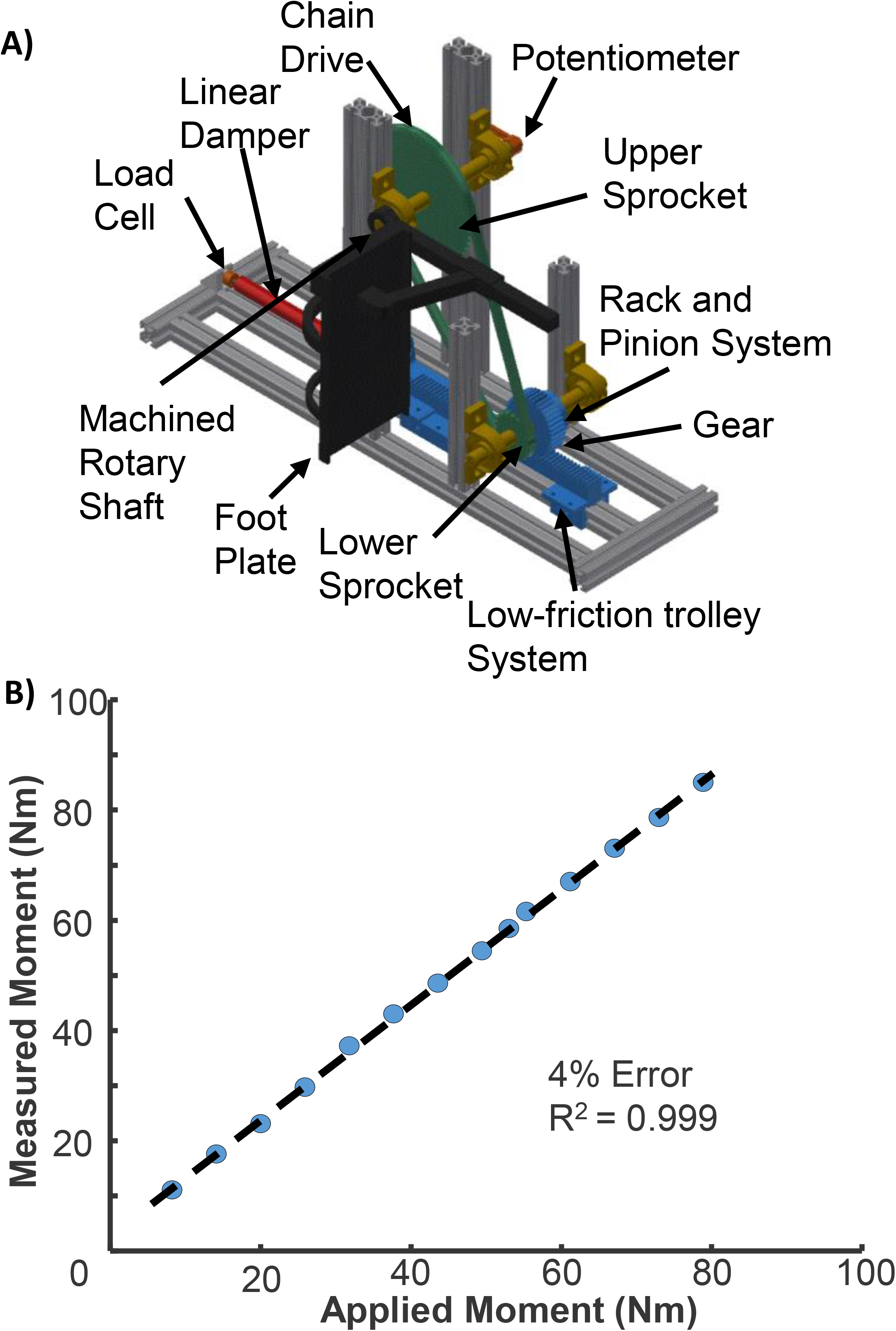
A) The isodamping dynamometer consists of three subsystems and an aluminum frame. The mechanical subsystem consists of a chain drive, rack and pinion system, and a linear damper. The arrangement and properties of each component dictates function of the machine as described in eq. 1 and 2. The subject interface system consists of a tapered spindle to accept commercial testing attachments and rubber stops to prescribe range of motion. The instrumentation subsystem consists of a potentiometer to measure angular position, a load cell to measure load transferred through the system, and a microprocessor to collect data. The aluminum frame can be designed for application specific considerations. For example, lower loads require less materials and bracing. B) Device measurements were validated by hanging weights at varying distances from an arm attached to the device.

To meet these requirements, we developed mechanical, subject interface, and instrumentation sub systems that we mounted to a custom-built aluminum frame (Supplemental Material). The *mechanical subsystem* transferred rotational loads about the ankle through a chain drive and a rack and pinion system to a linear damper (Easylift, Bansbach, Germany) that constrained rotational velocity as a function of the applied moment to the system. To ensure subject safety, we designed our isodamping dynamometer to act as passive energy sink, which constrains velocity by forcing incompressible oil through an orifice with an adjustable diameter. We mounted the linear damper to an aluminum frame using clevis pins, allowing any number of testing rotational velocities to be quickly changed by hot swapping dampers of different settings. We developed a mechanical model of the device to aid in its design and validation. Using this model, we derived the two following functions to predict device performance:

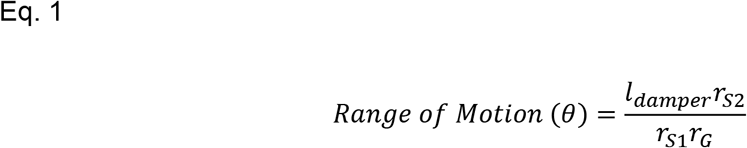

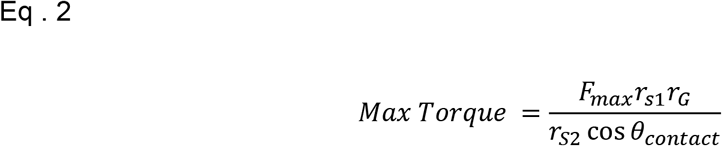

Where *l*_*damper*_ is the maximum displacement of the damper, *r*_*S*2_ is the radius of the lower sprocket in the chain drive, *r*_*S*1_ is the radius of the upper sprocket in the chain drive, *r*_*G*_ is the radius of gear in the rack and pinion system, *F*_*max*_ is the maximal force rating of the damper, and *θ*_*contact*_ is the contact angle between the gear and rack in the rack and pinion system. The *subject interface subsystem* consisted of all device components pertaining to subject testing and mobile deployment. One end of the upper spindle (8017T13, 1’’ Tapped D-Profile Rotary Shaft, McMaster-Carr, Robbinsville, NJ, USA) was machined to accept commercial dynamometer attachments. Range of motion was set by securing rubber washings in the track to limit the damper stroke. The *instrumentation subsystem* measured and logged the angular position, velocity, and moment about the ankle. We measured ankle angle using a simple rotary potentiometer (Linear 100k Ohm, Sparkfun, Niwot, CO, USA) and ankle moment using a load-cell (200 kg, TAS501, HTC-Sensor, Xi’an, China) in series with the linear damper coupled with a mechanical model of the system. We used a microcontroller (Metro Mini, Adafruit, New York, NY, USA) and a load cell amplifier (HX711, Sparkfun, Niwot, CO, USA) to condition the load-cell signal. The potentiometer and load-cell were both connected to a microcontroller (Arduino Mega REV 3, Arduino, Boston, MA, USA) that sampled data at 100 Hz and exported these data to a personal computer via serial port connection through a graphical user interface (MATLAB 2018a, The Mathworks, Natick, MA, USA).

### Mechanical Validation

We calibrated angular position and moment measurements prior to human subject testing. We applied known moment to the system by hanging weights at specific distances about the rotational shaft. To confirm the mechanical validity of our device, we correlated the applied moments with the experimentally measured moments from the device. This mechanical validation showed very strong agreement (R^2^ = 0.999, maximum of 4% full scale error) between the measured moments and the reference values (**Fig. 1B**).

### Human Subject Testing

Six healthy adults (3 male, 3 female, 30.2 ± 2.6 years, 25.5 ± 5.3 BMI) performed a series of plantar flexion contractions after providing written informed consent in this study approved by the University of Pennsylvania Institutional Review Board (824466). We assessed plantar flexor function on a commercially-available isokinetic dynamometer (System 4, Biodex, Shirley, NY, USA) and our novel isodamping dynamometer. We tested each subject resting prone on a treatment table with their right foot secured to the same foot attachment (Ankle Attachment for Biodex System 4, Biodex, Shirley, NY, USA). These subjects performed three plantar flexor contractions at three increasing effort levels (minimal, moderate, maximal) at three speed settings on both the isokinetic dynamometer (30, 120, and 210 degrees/s) and three damping settings on the isodamping dynamometer (soft, intermediate, and hard damping). To mitigate possible learning effects, we randomized the order of which dynamometer was tested first and the order of each of the test speeds. During pilot testing, we experimentally determined damping settings on the variable linear dampers that constrained rotational velocities within the range of speeds during isokinetic dynamometer testing. For each trial, we measured ankle angular positon, angular velocity, ankle moment and used these variables to calculate plantar flexion kinetics on each dynamometer. Prior to analysis we passed all data through a 6 Hz low-pass Butterworth filter.

We opted to have subjects perform contractions at variable effort levels to evaluate device performance across a wide range of mechanical demands. For each minimal effort trial, we verbally encouraged subjects to “rotate your ankle but don’t push.” For each moderate effort trial, we verbally encouraged subjects to “rotate your ankle but don’t push as hard as you can.” For each maximal effort trial, we verbally encouraged subjects to “push as hard and as fast as you can.” Following each trial, we provided verbal feedback of their peak moment after each contraction for reference (Drazan et al., 2019) and encouraged subjects to beat their previous peak moment during maximal effort contractions moment(McNair et al., 1996).

To determine the validity of isodamping dynamometry to quantify plantar flexion function, we assessed 1) similarities in peak plantar flexion moments between isokinetic and isodamping dynamometry and 2) velocity constraints imposed by isokinetic and isodamping dynamometry at different effort levels. We fit a quadratic curve (**Fig. 2**) to peak plantar flexion moment measurements (y-axis) at different speeds (x-axis) because previous studies demonstrated the force-velocity properties of muscle during dynamometry (Hauraix et al., 2017; Trappe et al., 2001).

**Fig. 2.**
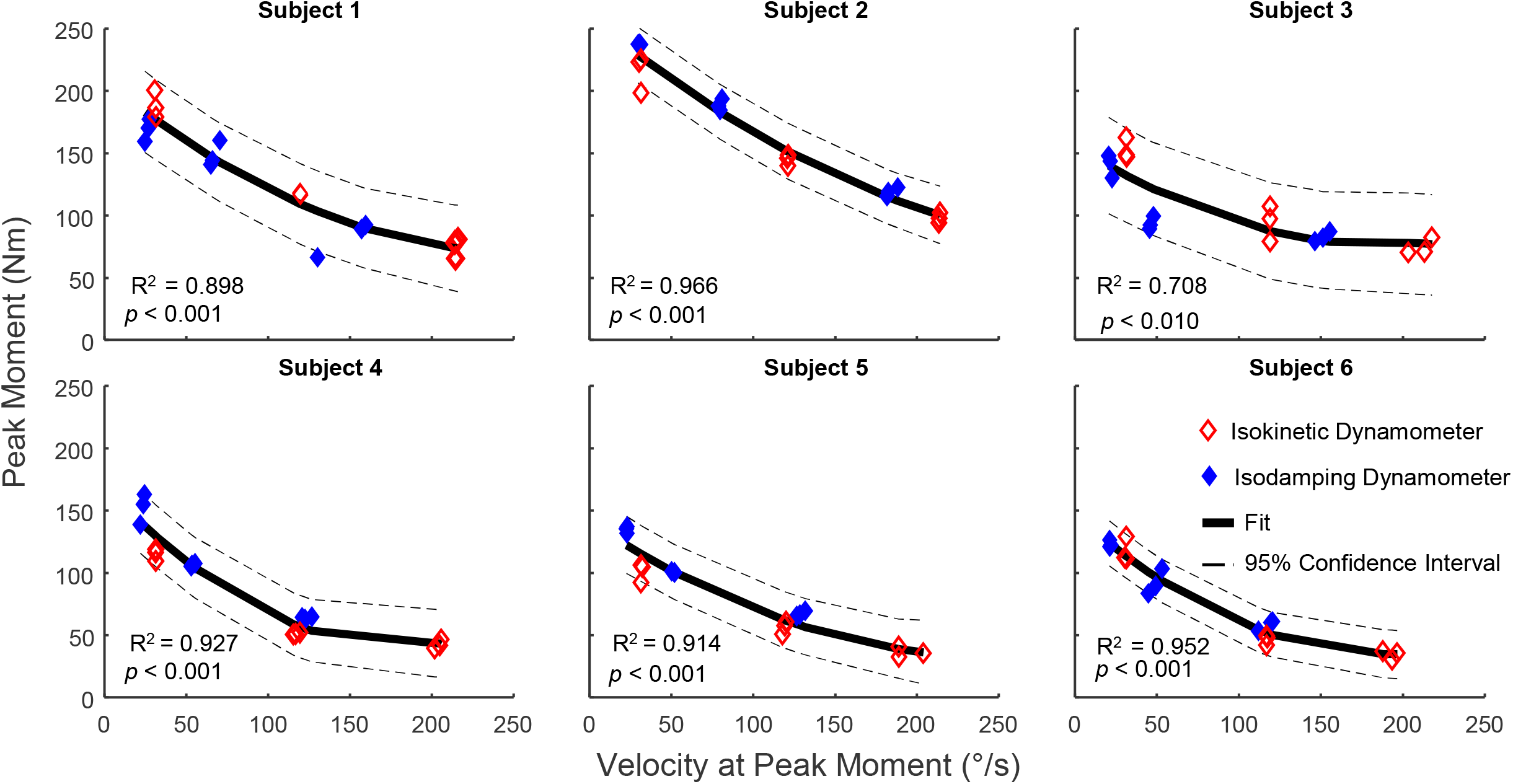
Measurements of peak moment and velocity at peak moment during maximal effort contractions fit on the same quadratic curve across both devices for all subjects (R^2^>0.7, *p* <0.01).

We considered the isodamping dynamometer valid if the isokinetic and isodamping dynamometer peak moment data fit on the same quadratic curve. We also established the load-velocity limitations of both isokinetic and isodamping dynamometry (**Fig. 3**). To evaluate this, we performed linear regression on velocity at peak moment and peak moment on all trials and subjects, grouped by speed condition.

**Fig. 3.**
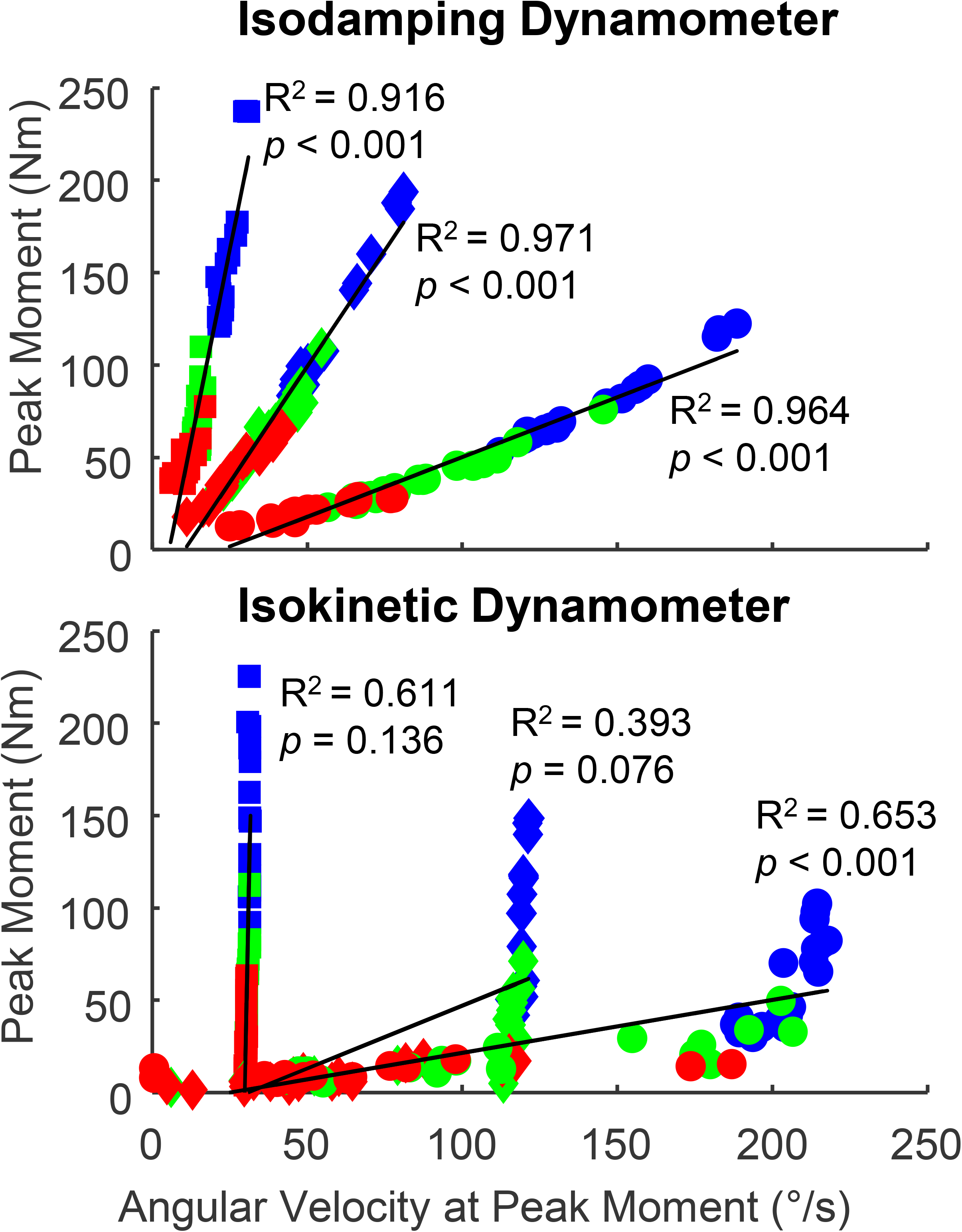
A) Measurements of peak moment and velocity at peak moment were strongly linearly correlated across all subjects and effort levels for the three damping settings on the isodamping dynamometer (R^2^>0.91, *p* <0.001). B) Measurements of peak moment and velocity at peak moment were not linearly correlated for the isokinetic dynamometry across subjects and effort levels except for the 210 °/s.

## RESULTS

Peak moment and velocity at peak moment was strongly correlated (R^2^ > 0.708, P ≤ 0.008) for all subjects during maximum effort contractions (**Fig. 2**). Across all effort levels and subjects, the isodamping dynamometer exhibited a strong, significant linear correlation (R^2^ > 0.91, P < 0.001, **Fig. 3A**). In contrast, the isokinetic dynamometer exhibited stepwise response where damping only occurred when the isokinetic threshold was reached (**Fig. 3B**). The linear relationship between peak moment and velocity at peak moment was only significant for the 210 °/s condition (R^2^ = 0.65, P < 0.001).

## DISCUSSION

We developed a mobile and low-cost alternative to traditional isokinetic dynamometers for assessing joint-level kinetics. Our isodamping dynamometer is significantly smaller (0.75 m^2^ footprint), lighter (30 kg), and lower cost (~2,200 dollars) compared to commercially-available isokinetic dynamometers (5.95 m^2^ footprint, 450 kg, and 40,000 dollars respectively)(Grabowski et al., 2017). Our damping approach limits velocity passively as a function of moment applied to the system rather than using an electrical motor to constrain velocity. Isodamping dynamometry also has the potential to monitor patient recovery in the clinic or field and increases the feasibility of collecting data from large subject cohorts. Isodamping dynamometry reproduces isokinetic dynamometers results at maximal effort across a cohort of six subjects (**Fig. 2**) and it also prescribes motion if maximal effort is not achieved (**Fig. 3**). While measurements of sub-maximal efforts are not typically used as clinical indicator or in research, it could have utility for reliably testing subjects who do not have the capacity to reach higher isokinetic speed (e.g. the elderly, infirm, or injured).

There are limitations to this device. While commercial isokinetic dynamometers are expensive, they have useful features such as being able to establish range of motion prior to testing and the ability to lock the ankle in place before the start of a trial. In contrast, with our device, the subject needs to voluntarily dorsiflex to hold their foot in place before a trial. In addition, our device has potential to be less expensive. We utilized a commercially-available ankle attachment for consistency between devices during testing. This ankle attachment was approximately 1/3 of the overall cost of the isodamping dynamometer. Designing a new lower-cost subject interface has potential to decrease the price. Additionally, while the new device closely matched the performance of the isokinetic dynamometer during maximal effort trials, the lowest damper setting was too difficult for the weaker subjects (**Fig. 2**: Subjects 4,5,6) reach high velocities (>140 °/s). Therefore, we recommend pre-selecting damper parameters based on the population being studied. Because isodamping dynamometry velocities constraints depend on the strength of the subject and damper parameters, it is not practical to prescribe specific speed constraints. Instead, we recommend testing joint kinetics at several different damper settings to define the force-velocity properties of the joint being tested (**Fig. 2**). By defining subject-specific force-velocity properties, isodamping dynamometry provides a more complete reflection of joint-level functional capacity than typical mobile measurement devices like handheld dynamometery while having a lower cost than traditional isokinetic dynamometers.

In summary, we developed and validated a novel isodamping dynamometer that provides a new opportunity to accurately assess joint-level kinetics. We developed this device to expand our clinical research footprint, which is currently limited by immobile and expensive testing systems. We believe that other researchers in biomechanics will also benefit from isodamping dynamometry because biomechanics research has been hampered by the widespread use of small, unrepresentative convenience samples (Vagenas et al., 2018). This may be in part due to limited options for mobile data collection platforms (Mavroidis et al., 2005; Verheul et al., 2020). This new isodamping dynamometer device has potential to allow researchers to collect data outside of the laboratory setting. For example, large data collections could be performed outside the lab is during community outreach events (Drazan, 2019; Shultz et al., 2019). Finally, the isodamping dynamometer has a tapered spindle that is compatible with commercial dynamometer attachments can assess other joints including the knee, the hip, or upper body with minor modifications to the frame system.

## Supporting information

Parts list for dynamometer

## ACKNOWLEDGEMENTS

This research was supported by NIH grant# K12GM081259.

